# Elucidating the crosstalk between inflammation and DNA damage pathways in the pancreatic beta-cell through the diabetes susceptibility gene, *TCF19*

**DOI:** 10.1101/2021.04.06.438736

**Authors:** Grace H. Yang, Danielle A. Fontaine, Sukanya Lodh, Joseph T. Blumer, Avtar S. Roopra, Dawn Belt Davis

## Abstract

*TCF19* is a gene that is associated with both type 1 diabetes (T1DM) and type 2 diabetes (T2DM) in genome-wide association studies. Prior studies have demonstrated that TCF19 knockdown impairs β-cell proliferation and increases apoptosis. However, little is known about its role in diabetes pathogenesis or the effects of TCF19 gain-of-function. The aim of this study was to examine the impact of TCF19 overexpression in INS-1 β-cells on proliferation and gene expression. With TCF19 overexpression, there was an increase in nucleotide incorporation without any change in cell cycle gene expression, alluding to an alternate process of nucleotide incorporation. Analysis of RNAseq of TCF19 overexpressing cells revealed increased expression of several DNA damage response (DDR) genes, as well as a tightly linked set of genes involved in cell stress, immune system processes, and inflammation. This connectivity between DNA damage and inflammatory gene expression has not been well studied in the β-cell, and suggests a novel role for TCF19 in regulating these pathways. Future studies determining how TCF19 may modulate these pathways may provide potential targets for β-cell survival.

## 1. Introduction

The pancreatic β-cell is susceptible to many different stressors including oxidative stress, endoplasmic reticulum (ER) stress, and inflammation^1,2^. These stressors are exacerbated in patients with obesity, insulin resistance, and diabetes ^3–5^. This can lead to β-cell apoptosis and reduced β-cell mass. Pancreatic islets from patients with T2DM have increased ER stress which can lead to β-cell dysfunction and apoptosis^6,7^. In addition, increased circulating cytokines and localized islet inflammation are characteristics of T2DM patients and can contribute to β-cell death^8^. Hyperglycemia, as well as metabolic abnormalities associated with diabetes can lead to oxidative stress, resulting in increased intracellular reactive oxygen species (ROS) that contribute to β-cell dysfunction ^9,10^. While many of these sources of β-cell stress have been well studied, there are other factors that can lead to β-cell dysfunction and apoptosis that have received little attention. In particular, DNA damage has started gaining attention in recent years as having a role in diabetes pathogenesis. The diabetic microenvironment in the islet presents oxidative stress and inflammatory insults that have been shown to increase DNA damage in the islet ^11–14^. Additionally, DNA damage to islets elicited by the β-cell toxin, streptozotocin (STZ), causes an elevation of proinflammatory cytokines^12^. However, this inflammatory response is attenuated after inactivation of the master DNA repair gene, ataxia telangiectasia mutated (ATM) ^12^. Horwitz et al. also demonstrated that the β-cell DDR was more frequent in islets infiltrated by CD45+ immune cells^12^. This brings to light a fascinating connection between DNA damage and inflammatory responses in the islet. A better understanding of the intersection between these two processes will provide potential regulatory targets to reduce and resolve DNA damage and inflammatory stress on the β-cell that may serve to help maintain adequate β-cell mass and function in diabetes.

In humans, the gene *TCF19* (transcription factor 19) is associated with both T1DM and T2DM in genome-wide association studies^15–18^. *TCF19* is expressed in human islets and shows a positive correlation with BMI in nondiabetic subjects^19^. In mice, *Tcf19* is ubiquitously expressed; however, its expression is highest in the pancreatic islet and increases with obesity when β-cells are known to increase proliferation ^19^. Others have similarly identified *Tcf19* as a gene upregulated in proliferating β-cells and found that knockdown of Tcf19 impairs insulin secretion in a human β-cell line^20–24^. We have previously demonstrated that siRNA-mediated knockdown of *Tcf19* in rat insulinoma INS-1 cells reduces β-cell proliferation and survival and impairs cell cycle progression beyond the G1/S checkpoint ^19^. Additionally, *Tcf19* knockdown increases apoptosis via reduced expression of genes involved in the maintenance of ER homeostasis and increased expression of proapoptotic genes^19^.

The TCF19 protein contains a forkhead association (FHA) domain, which is a phosphopeptide recognition domain commonly found in many transcription factors that participate in DNA repair and cell cycle regulation ^25^. The human TCF19 (hsTCF19) protein, but not the mouse protein, also harbors a plant homeodomain (PHD) finger, allowing it to interact with chromatin. PHD finger proteins are often considered “chromatin readers” that recognize modified histones and can recruit additional transcriptional machinery to these areas ^26^. Specifically, the tryptophan residue at position 316 in hsTCF19 has been shown to bind to chromatin via tri-methylated histone H3 and to regulate cell proliferation in liver cells via this interaction^27^. Taken together, these characteristics support the role of TCF19 as a transcriptional regulator of β-cell proliferation and survival.

The purpose of this study was to determine the effect of TCF19 overexpression on proliferation and survival in the β-cell. We find that TCF19 is regulating a node of tightly interconnected genes that have roles in cell stress, inflammation, and antiviral responses. Additionally, we find that TCF19 overexpression leads to significant upregulation of several DDR genes. In this study, we overexpressed TCF19 in INS-1 cells and found that TCF19 overexpression does not induce proliferation or cell cycle progression. Rather, there was significant upregulation of a tightly interconnected set of genes involved in inflammation, antiviral, immune system, and DDR pathways, alluding to a previously unexplored role for TCF19 in the β-cell. Using a novel analysis for potential transcriptional co-regulators of these upregulated genes, we identified STAT1, STAT2, and IRF1 as likely drivers of the tight transcriptional gene network. Interestingly, there was no measurable activation of these transcription factors, indicating alternative means of regulating the inflammatory and DNA damage gene expression. These findings not only identify an intriguing connection between DNA damage and inflammatory responses in the β-cell, but elucidate a novel role for TCF19 in modulating these two pathways.

## 2. Results

### 2.1 Human TCF19 overexpression increases H-thymidine incorporation in INS-1 cells but does not change cell cycle gene expression

Based on our original studies on TCF19, we concluded that TCF19 was necessary for normal β-cell proliferation, as TCF19 knockdown led to impaired cell cycle progression, reduced ^3^H-thymidine incorporation, and G1/S cell cycle arrest^19^. We next wanted to determine if increased levels of TCF19 could drive β-cell proliferation, and therefore, we overexpressed hsTCF19 in INS-1 rat insulinoma cells. The human TCF19 protein was chosen for overexpression as it contains the PHD finger domain which is known to mediate interactions with methylated histones (specifically trimethylated histone 3 at lysine 4 (H3K4me3)), and this domain is not found in the rodent protein^27,28^. As we have not yet identified a reliable and specific TCF19 antibody, we generated a C-terminal myc-tagged TCF19 to allow for probing on the western blot. TCF19 overexpression was confirmed at both the mRNA and protein level (**Fig 1A).** As an assay to assess proliferation, we measured ^3^H-thymidine nucleotide incorporation in cells expressing hsTCF19 vs. empty vector control. INS-1 cells overexpressing hsTCF19 showed a significant two-fold increase in ^3^H-thymidine nucleotide incorporation suggesting increased cell proliferation **(Fig 1B**). To confirm that the ^3^H-thymidine nucleotide incorporation observed correlated with an increase in the expression of cell cycle genes, as would be expected in a dividing cell, we assessed cell cycle gene expression with qRT-PCR. Interestingly, there was no significant change in expression of cell cycle genes **(Fig 1C**). In addition, there was no significant change in levels of the proliferative marker, Ki67 (**Fig 1C**). We concluded that overexpression of hsTCF19 in INS-1 cells does not lead to transcriptional activation of cell cycle genes, suggesting an alternate process for nucleotide incorporation that does not result in cell cycle progression. DNA repair may be an alternative pathway that leads to increased ^3^H-thymidine nucleotide incorporation^28^. DNA damage and repair responses are important in preserving genome integrity, and an accumulation of DNA damage without sufficient repair can result in cell cycle arrest at the G1/S checkpoint. However, qRT-PCR (**Fig 1C**) showed no significant change in cell cycle inhibitors *Cdkn2c* (*p18*), *Cdkn1a* (*p21*), and *Cdkn1b* (*p27*) with hsTCF19 overexpression, suggesting that there wasn’t necessarily any induction of DNA damage leading to cell cycle checkpoint arrest. We next hypothesized that if hsTCF19 overexpression is affecting DNA repair, it may elicit a change in cell viability. However, after staining cells with trypan blue, we found that the percentage of live cells was not significantly affected by hsTCF19 overexpression (**Fig 1D).**

**Fig 1.**
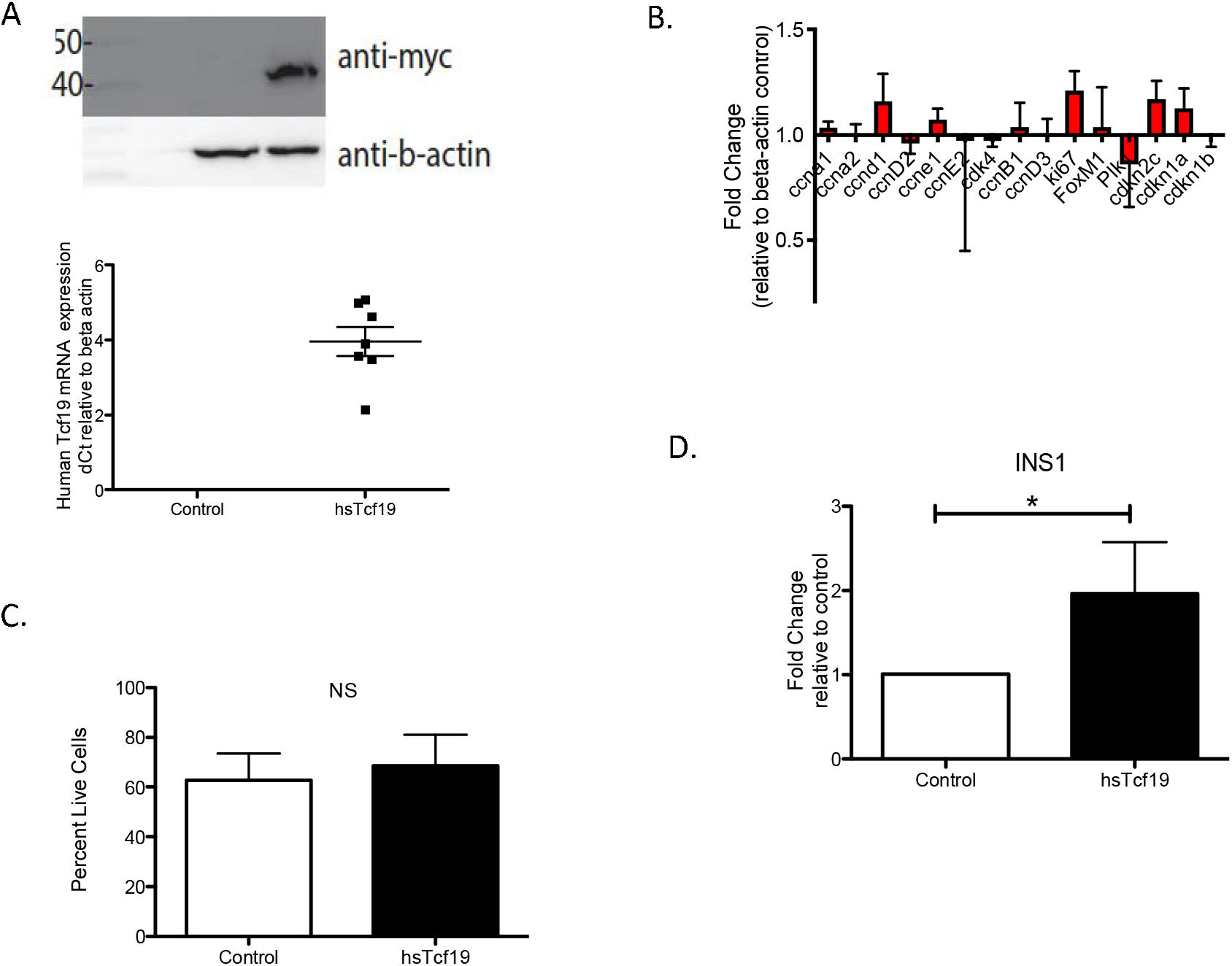
(A) Overexpression of human TCF19 in INS1 cells was confirmed by real time PCR and western blot. (B) hsTCF19 overexpression does not lead to any significant changes in cell cycle gene expression (n=5) (C) Overexpression of hsTcf19 in INS1 does not affect cell viability (n=5) (D) Overexpression of hsTCF19 in INS1 leads to increased tritiated thymidine incorporation (n=5). Data are means ± SEM *P<0.05.

### 2.2 RNA-Seq analysis reveals a role for TCF19 in regulating viral, inflammatory, and DNA damage genes

To obtain a more global perspective on what genes TCF19 could be regulating, we performed RNA-seq analysis of INS-1 cells overexpressing hsTCF19. Notably, this revealed only a relatively small number of differentially expressed genes. Of the 16C genes differentially expressed between the groups (false discovery rate <0.05), 136 genes were upregulated and 24 were downregulated (**Supplemental Table 1**), suggesting that TCF19 likely acts as a positive regulator of transcription. Overrepresentation test with PANTHER on the upregulated gene set revealed an enrichment for pathways including the interferon signaling response, immune system processes, and response to viruses (**Fig 2A**).

**Fig 2.**
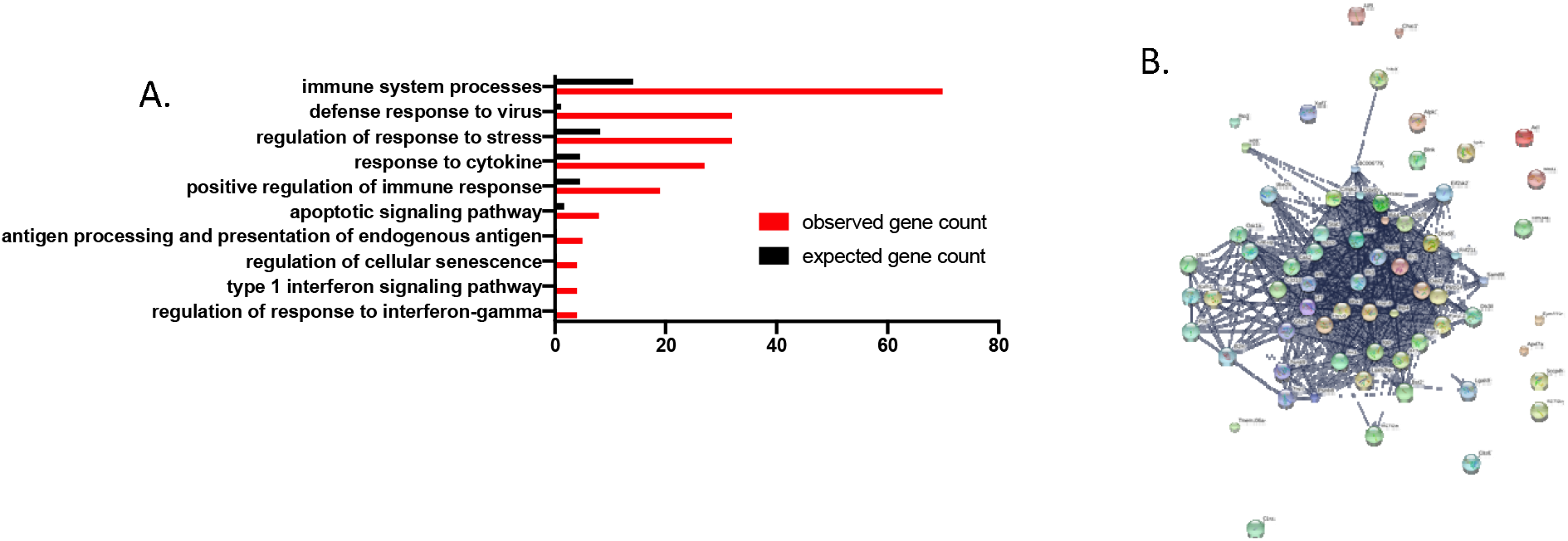
RNA seq analysis of INS1 cells overexpressing hsTCF19 identifies upgregulated genes that form a tight node of interconnected genes with roles in viral and stress response pathways (A) PANTHER overrepresentation GO terms test shows upregulation of genes involved in viral response and response to cytokines (fold change > 1.5, FDR<0.05) (B) STRING analysis on DE upregulated gene set shows tightly interconnected network of genes.

To determine the relationship between the significantly upregulated genes from our RNAseq data, we performed STRING analysis. STRING pulls from numerous sources to predict potential protein-protein interactions to assess for clusters of proteins that may have functional similarity or similar co-expression^23^. STRING analysis on the upregulated gene set revealed a tight connection between almost all upregulated genes, suggesting that TCF19 may be regulating one cluster of interconnected genes (**Fig 2B**). We hypothesized that this cluster of genes may have roles in viral and interferon responses, as well as the DDR.

Among the upregulated genes, several are known to be involved in DDR and repair pathways (*Parp9, Parp10, Parp12, Parp14*)^29^. In particular, *Parp9* and another gene from our dataset, *Dtx3l*, have been shown to work as a complex to promote DNA repair^30,31^. Other significantly upregulated genes included those from the oligoadenylate synthase (Oas) family (*Oas1i, Oasl2, Oas2, Oas1a, Oas1g, Oas1f*),> which are stimulated by type 1 interferons in response to viral infections. However, they can also be activated by DNA damage, where they may have roles in Poly(ADP-ribose) (PAR) synthesis and interacting with PARP1 during DNA repair^32,33^. *Mx1, Ddx60* and *Usp18*, are genes with known antiviral roles and were also significantly upregulated ^34–36^ (Supplementary Table 1). These observations suggest that TCF19 may play a previously unreported role in the DNA damage response and viral and inflammatory response pathways.

To assess the extent to which the findings in this overexpression model could be translated to human islets, we overexpressed hsTCF19 in human islets and assessed several of the differentially expressed genes from the RNAseq dataset in INS-1 cells (**Fig 3A and B).** Notably, DNA damage transcript levels for genes *PARP9* and *DTX3L* were significantly upregulated in human islets overexpressing hsTCF19 compared to the empty vector control islets (**Fig 3B).** Antiviral genes *MX1* and *DDX60* were also significantly upregulated.

**Fig. 3.**
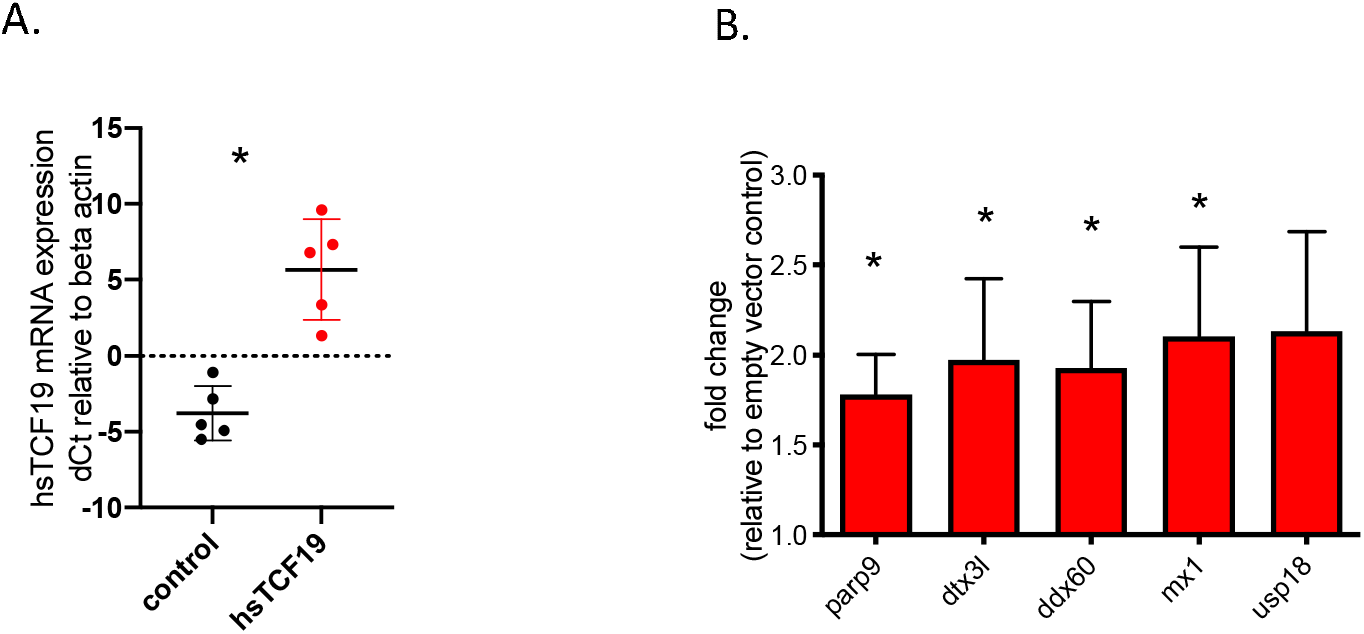
Human TCF19 overexpression in human islets upregulates key DNA damage repair genes as well as interferon response genes (A) TCF19 overexpression in human islets was confirmed with qRT-PCR. (B) TCF19 overexpression in human islets leads to upregulation of genes (qRT-PCR) that were also upregulated in INS-1 cells (n=5). Data are means ± SEM *P<0.05.

### 2.3 Mining Algorithm for GenetIc Controllers (MAGIC) Analysis for common transcriptional regulators

To look for common transcriptional regulators associated with the promoters of the upregulated RNA-Seq gene set, we performed MAGIC analysis^37^. These analyses, which are based upon annotated databases including ENCODE ChIP-seq data, revealed Signal Transducer and Activator of Transcription (STAT)1 and STAT2 as positive drivers of this gene set (**Fig 4**). The associations were striking with p-values of 7.81E-19 and 3.23E-20, respectively. Specifically, STAT1 and STAT2 are known to interact with the promoter of 17 genes in our dataset. However, these associations in ENCODE were not determined in β-cells or islets and were often based on experiments involving interferon stimulation. There was strong enrichment of these genes compared to overall promoter interactions for these STAT proteins across the genome, suggesting that TCF19 leads to upregulation of genes that can also be regulated by the STAT proteins. Additionally, Interferon Response Factor (IRF)1, a transcription factor important in both innate and adaptive immunity, also showed striking enrichment for promoter interactions with the upregulated gene set (p = 2.77E-16). As these transcription factors could be potential regulators of the upregulated genes in our dataset, we assessed for activation of these transcription factors. However, densitometric quantification of protein levels in cells overexpressing TCF19 showed only a modest increase in active phosphorylated STAT1 (**Fig 5A**). There was no change in IRF1 levels (**Fig 5C**). STAT2 was not detectable in the INS-1 cells. Taken together, this suggests that TCF19 does not directly modulate the levels or phosphorylation status of STAT1, STAT2, or IRF1 in β-cells.

**Fig. 4.**
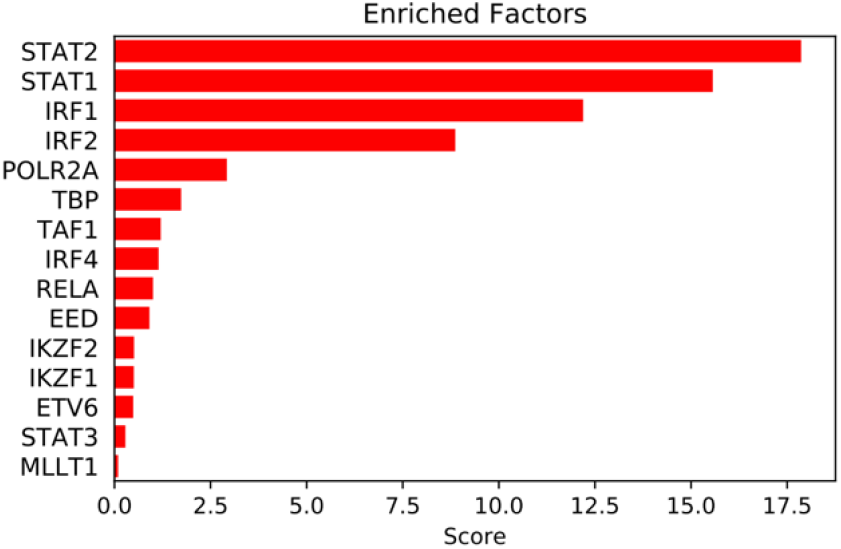
MAGIC analysis on the list of upregulated genes after TCF19 expression in INS-1 cells identifies significant enrichment for genes with known ChIP signals for STAT1, STAT2, and IRF1 in their promoters. Genes used in analysis were those that were upregulated more than 2-fold with an associated FDR < 5%.

**Fig. 5.**
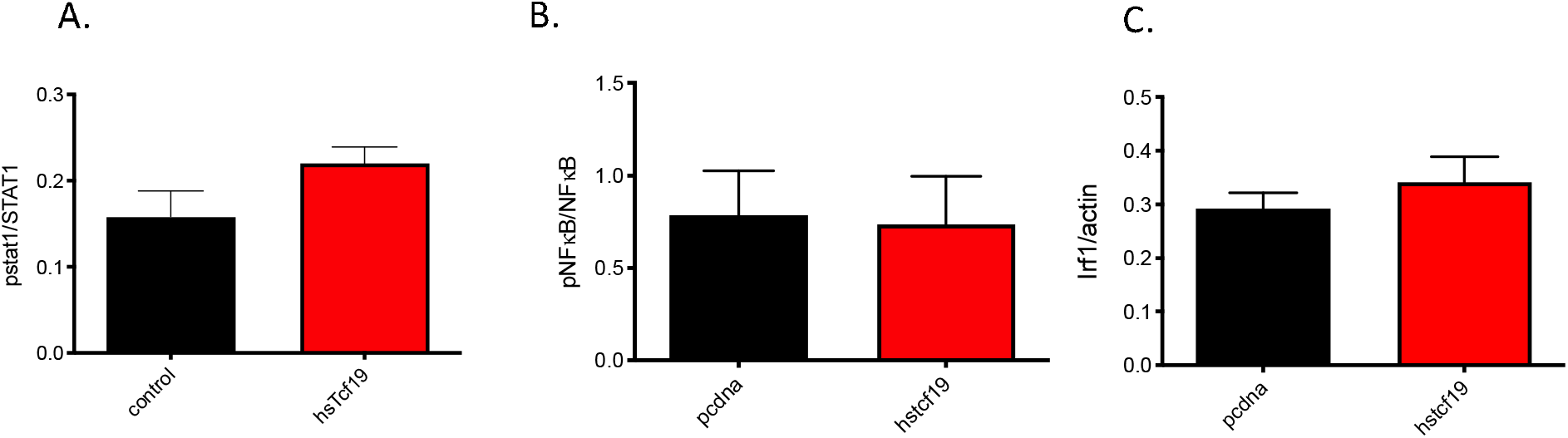
Overexpression of TCF19 in INS1 cells does not lead to increased activation of transcription factor targets. Western blot quantification of (A) phospho-STAT1/STAT1 protein expression does not show a statistically significant difference between control and hsTCF19 overexpressing cells. (B) There is also no difference in phosphoNF-κB, / NF-κB levels with hsTCF19 overexpression (C) IRF1 protein levels are not significantly different (n=5). Data are means ± SEM.

Although not identified as a potential co-regulator in MAGIC analysis, Nuclear Factor Kappa-B (NF-κB) has a well characterized role in mediating inflammation and is also activated by the cGAS-STING pathway, which is a component of the innate immune system that functions to detect cytosolic DNA and leads to the production of type 1 interferons^38^. Therefore, we predicted that NF-κB may be a possible regulator of the upregulated gene set. However, we found no increase in the phosphorylation of NF-κB (**Fig.5B**) with Tcf19 overexpression.

## Discussion

Inflammation is a pathophysiological state associated with both T1DM and T2DM. In T1DM, immune cells are critical mediators of islet inflammation through their secretion of cytokines such as interleukin 1 beta (IL-1beta) and tumor necrosis factor-alpha (TNF-alpha)^39^. Additionally, there is substantial evidence suggesting that triggering events such as a viral infection may initiate the β-cell damaging process in a large proportion of the patients ^40, 41^. In T2DM, obesity induces chronic, low grade inflammation which leads to the activation of inflammatory pathways and the release of pro-inflammatory cytokines and adipokines ^38,42,43^. Inflammation not only exacerbates insulin resistance and promotes β-cell death, but can also contribute to DNA damage^15^.

In this study, we describe a role for the diabetes susceptibility gene, *TCF19*, in the inflammatory and DNA damage pathways. We find that TCF19 overexpression significantly increases expression of inflammatory and DDR genes, suggesting a novel role for TCF19 in regulating these two pathways. We find that the significantly upregulated genes from TCF19 overexpression are tightly associated, and we describe potential transcription factor co-regulators of these genes. This brings to light an interesting crosstalk between the inflammatory and DNA damage pathways in the β-cell.

Knockdown of *Tcf19* has been shown to result in cell cycle arrest. While we show here that overexpression of hsTCF19 does not result in significant changes in cell cycle genes, hsTCF19 overexpression results in increased expression of DDR genes. The DDR is made up of DNA damage sensing proteins, transducers, and effectors. Once an aberrant DNA structure is recognized, downstream phosphorylation cascades within the DDR network are initiated with many of the downstream effector proteins having roles in promoting cell cycle arrest ^44^. This allows time for the cell to repair the damaged DNA. Other effector proteins upregulate DNA damage repair genes or promote senescence or apoptosis in the face of unrepairable DNA damage ^40^. With TCF19 overexpression, we find an increase in genes involved in the DDR but no decrease in cell viability, suggesting that these cells are not undergoing apoptosis. Additionally, the lack of significant change in cell cycle genes including cell cycle inhibitors suggests there is no DNA damage-induced cell-cycle arrest. Notably, these experiments were all performed in the absence of any inducers of DNA damage or interferons, yet we observed upregulation of classic interferon-response genes. Therefore, enhanced TCF19 expression alone is sufficient to independently activate these pathways. Since overexpression of TCF19 led to upregulation of many DNA damage repair genes, this suggests that within the DDR network, TCF19 most likely plays a role as a transcriptional regulator that may promote DNA damage repair.

TCF19 knockdown leads to cell cycle arrest at the G1/S transition^19^. This is consistent with the cell cycle arrest that occurs upon DNA damage to cells in the G1 phase to prevent entry into the S phase ^45^. Sustained DNA damage can eventually result in cellular apoptosis ^46^. We previously showed that 3-7 days of *Tcf19* knockdown led to an increase in cells undergoing apoptosis and a decrease in cell viability. Combining these prior results with current data, we propose that cells lacking TCF19 are inefficient at repairing DNA damage, ultimately leading to cell cycle arrest or cell death due to accumulated DNA damage. We hypothesize that with TCF19 overexpression, however, DNA damage repair is upregulated.

Interestingly, many of the genes upregulated by TCF19 overexpression are also involved in interferon and immune responses. Additionally, Gene Ontology analysis revealed an overrepresentation of genes involved in viral response signaling. This signature of viral, inflammatory and DNA damage responses brings to light an interesting and emerging field regarding the connection between DNA damage and the interferon response. Treatment of cells with etoposide, an agent that induces double stranded DNA breaks, leads to the induction of interferon (IFN) stimulated genes regulated by NF-κB^47^. The cGAS-STING pathway is a component of the innate immune system which functions to detect cytosolic DNA and activates the Stimulator of Interferon Genes (STING), resulting in the production of type 1 interferons ^48^. However, after etoposide treatment, there is noncanonical activation of the STING pathway by the DNA repair proteins, ATM and PARP1^48^. Additionally, the DNA sensor cyclic GMP-AMP synthase (cGAS) has been shown to be shuttled to the nucleus under conditions of DNA damage ^49^. Given these connections, we also looked for an increase in phospho-STING after TCF19 overexpression but did not see any significant changes (data not shown). Further exploration of possible connections between the cGAS-STING pathway and DNA damage and inflammatory responses in the β-cell remain as intriguing new directions for future study.

While these studies show that DNA damage can lead to inflammatory gene expression, inflammation can also induce DNA damage. Chronic inflammation can lead to the production of ROS which are capable of DNA damage through the formation of free radicals and DNA lesions^50^. To further support the coordinate regulation between these two pathways, it has been shown that viruses can activate the DDR network and also inhibit several DDR proteins ^51^. As viral infection is an important initiating factor in T1DM, this could serve as a potential link between the two pathways where the immune system’s viral response may trigger DNA damage and progression to T1DM. It is likely that the DDR and inflammatory pathways are part of a positive feedback loop^52^. We see a dual response gene signature of viral/interferon and DNA damage processes with TCF19 overexpression, suggesting that TCF19 may regulate both of these processes. However, we also acknowledge the possibility that TCF19 may regulate just one of these processes, and in turn, be indirectly affecting the other.

STRING analysis further supports the tight association between the DNA damage and inflammatory genes in our data set. MAGIC analysis revealed STAT1, STAT2, and IRF1 as common regulators of this gene set. These transcription factors have well characterized roles in response to interleukins and interferons, specifically type 1 interferons^53^. However, there have also been studies showing a role for these transcription factors in the DDR and repair pathway^51^. A few of these transcription factors have been found to be responsible for the induction of interferon alpha and gamma genes in response to DNA damage or have roles in regulating DNA damage repair proteins^54^.

While we did not observe direct increases in phosphorylation or protein levels of these transcription factors, phosphorylation events can be transient and tightly regulated. It is possible that the time point of harvest (48 hours post transfection) may have been too late to capture the phosphorylation event. Additionally, while we chose to look at phosphorylation events for activation of these transcription factors, other types of post translational modifications, such as those that may work to alter chromatin structure or recruit histone modifiers cannot be ruled out. Notably, TCF19 has been shown to interact with histone 3 lysine 4 through its PHD finger to repress gluconeogenic gene expression and to modulate proliferation in HepG2 cells ^27,55^. Therefore, it is likely that TCF19 is not directly activating these transcription factors through phosphorylation events, but instead may bind to H3K4me3 at a transcriptionally active promoter and thereby impacts transcriptional activation. Additionally, the TCF19 protein harbors a FHA domain, which may allow binding to phosphor-epitopes on proteins ^56,57^. FHA domains are often found in proteins that are critical in the cell cycle and regulated through phosphorylation events, but are also found in proteins that are involved in the DDR^56^. The FHA domain of TCF19 contains a serine residue at position 78 (Ser78) that has been shown to be phosphorylated after DNA damage^58^. Ser78 in TCF19 is located within a Ser-Gln motif, which is recognized by kinases involved in the DDR such as ATM, and ATR. Therefore, it is possible that TCF19 is a downstream target of ATM or can alter gene expression by acting as a co-regulator to other kinases or recruiting phosphorylated transcriptional regulators to gene promoters.

Overall, we hypothesize that TCF19 is affecting DDR gene expression through histone modifications via the PHD finger and/or acts as a co-activator to DNA damage proteins by recruiting other DNA damage transcription factors to areas of active chromatin. While we did not directly measure an interaction of any of the transcription factors from the MAGIC analysis with relevant promoters in response to TCF19 overexpression, our data suggest that TCF19 either modulates their ability to activate transcription or may in fact simply be regulating the expression of these genes independently of these transcription factors.

The exact mechanism of how TCF19 modulates these inflammatory and DDR genes to promote diabetes susceptibility requires further investigation. Overall, our work highlights the complexity of regulation of gene expression involved in DNA damage and inflammatory response genes and alludes to the interesting crosstalk between these processes in the context of a diabetes susceptibility gene, *TCF19*. With respect to diabetes susceptibility, individuals with genetic variants of *TCF19* may be unable to properly regulate β-cell responses to DNA damage and inflammatory insults, therefore predisposing them to increased β-cell death. Future experiments will explore the nature by which TCF19 modulates DNA damage repair and inflammatory genes under conditions of stress. Furthermore, this will provide for potential therapeutic targets to prevent or attenuate DNA damage and inflammation to preserve functioning β-cells in at risk individuals.

## 4. Methods

### 4.1 Human islets and INS-1 cell culture

INS-1 E rat insulinoma cells were cultured in RPMI 1640 supplemented with 1% antibiotic-antimycotic (Gibco, 15240 – 062), 1% L-glutamine, 1% sodium pyruvate, and 10% fetal bovine serum. 2-Mercaptoethanol was added to a final concentration of 50uM to supplemented media before each use. Human islets were obtained from nondiabetic organ donors through the Integrated Islet Distribution Program. An exemption was granted for human islet work by the Institutional Review Board at the University of Wisconsin. Human islets were cultured in uncoated petri dishes with RPMI 1640 containing 8mM glucose, 10% heat-inactivated fetal bovine serum, and 1% penicillin/streptomycin. INS-1 cells and islets were cultured at 37°C and 5% CO_2_ in a humidified atmosphere.

### 4.2 Creation of TCF19 overexpression vector

The human TCF19 clone HsCD00002769 was purchased in the pDNR-Dual vector backbone (DNASU Plasmid repository). The pcDNA4-TO-myc/his B backbone vector (Invitrogen) was chosen for overexpression. This vector utilizes a CMV promoter, which ensures robust expression of the inserted gene of interest. Following the inserted Tcf19 sequence is both a C-terminal c-myc tag as well as six histidine residues to allow for identification of the overexpressed protein in the absence of reliable Tcf19 antibodies. The hsTCF19-pcDNA4 vector was created with In-Fusion HD cloning (Clontech) following kit instructions. Colonies were screened with PCR for insert size and then sequenced to confirm TCF19 insertions and sequence integrity.

### 4.3 Transfection with hsTCF19-his/myc-pcDNA4

INS-1 cells and islets were transfected with either hsTCF19 or pcDNA4 control, using Lipofectamine 2000 (Invitrogen). INS-1 cells were trypsinized and resuspended in transfection medium (RPMI 1640 supplemented with 1% L-glutamine, 1% sodium pyruvate, and 10% fetal bovine serum). Cells in transfection medium were then added to a hsTCF19 or control plasmid-Lipofectamine mixture at 2-5 μg DNA/5×10^6^ cells and plated. Transfection medium was removed 12–18 h post-transfection and replaced with complete growth medium. These conditions were the same for all INS-1 overexpression studies, including RNA-Seq sample preparation.

Human islets were washed in 1x PBS and resuspended in Accutase (Sigma) dissociation solution for 3 minutes at 37°C, with tube inversions every 30 seconds. Islets were then resuspended in 2mL transfection medium and plated into dishes. hsTCF19 or control plasmid-Lipofectamine 2000 mixture was added at 2ug DNA/1000 islets. Transfection medium was removed 12–18 h post-transfection and replaced with complete growth media.

### 4.4 Western blotting

INS-1 cells were harvested 48 hours after transfection and washed in ice-cold PBS. Cells were lysed in protein lysis buffer (0.05M HEPES, 1% NP-40, 2mM activated sodium orthovanadate, 0.1 M sodium fluoride, 0.01 M sodium pyrophosphate, 4mM PMSF, 1mM leupeptin, 2uM okadaic acid and Sigma Protease inhibitor cocktail). Cells were incubated in the lysis buffer on ice for 15 minutes with vortexing every 5 minutes. The protein concentrations were determined using Bradford protein assay. The protein samples were run on 4–10% SDS-PAGE gel and transferred to polyvinylidene difluoride (PVDF) membrane. Membranes were blocked in 5% milk in Tris-buffered saline with 0.1% Tween 20 (TBST) for 1 h at room temperature and were incubated overnight in primary antibody, washed 3X in TBST and incubated 1 h in secondary antibody. Blots were developed with Pierce ECL Western Blotting Substrate (Thermo Fisher) or Supersignal West Femto Maximum Sensitivity Substrate (Thermo Fisher), imaged with a GE ImageQuant charge-coupled device camera, and then quantified by densitometry with Image J 1.44o (http://imagej.nih.gov/ii). Primary antibodies and dilutions were as follows: Myc antibody (9E10:sc-40, Santa Cruz Biotechnology, 1:1000), Beta actin (8H10D10, Cell Signal, 1:1000), phosphorylated STAT1 Y701 (58D6, Cell Signaling, 1:1000), phospho-NF-κB p65 (93H1, Cell Signaling 1:1000), IRF1 (D5E4, Cell Signaling, 1:1000) all in 5% BSA-TBST.

### 4.5 Quantitative real-time PCR

RNA was isolated from INS-1 and human islets 48 hours post transfection using RNeasy cleanup kit (Qiagen) according to manufacturer’s protocol. Concentration and purity of RNA was determined using a NanoDrop ND-2000c Spectrophotometer, and 100-250 ng of RNA was reverse transcribed to make cDNA with Applied Biosystem High Capacity cDNA synthesis kit. Quantitative real-time PCR (qRT-PCR) reactions were carried out using Power SYBR green PCR Master Mix (Applied Biosystems) and the StepOnePlus Real Time PCR System (Applied Biosystems). Reverse transcriptase free samples were used as negative controls. All samples were run in triplicates with Cycle threshold (*Ct*) values normalized to ß-actin to yield ΔC*t*. Fold changes were then calculated between experimental and control samples: fold change 2^(ΔC*t*experimental - ΔC*t*ontrol)^. For gene expression in INS-1 cells, results were analyzed by nonpaired *t*-test of the ΔC*t* values, while human islets were analyzed by paired t-test. Significance was determined by *P*< 0.05. Primer sequences used are in Supplementary Table 2.

### 4.6 Viability

Transiently transfected INS-1 cells were harvested at 48 hours post-transfection by using a cell scraper to dislodge all cells and 10ul of cells were collected from each well. Cell viability was determined using trypan blue (Corning) staining using the TC-10 Automated Cell Counter (BioRad). Comparisons were made by paired *t*-test, including all technical and biological replicates; statistical significance was determined by *P*< 0.05.

### 4.7 Proliferation/^3^H-thymidine incorporation

To measure cell proliferation, transiently transfected INS-1 cells were incubated with ^3^H-thymidine (Perkin Elmer) at a final concentration of 1 Ci/ml in supplemented RPMI for 4 h. Cells were then trypsinized and washed three times with ice-cold PBS. DNA and protein were precipitated by the addition of ice-cold 10% trichloroacetic acid (TCA) and incubated for 30 min on ice. The precipitate was then pelleted at 18,000 *g* for 10 min at 4°C. Pelleted precipitate was solubilized in 0.3 N NaOH and vortexed for 15 min. Radioactivity was measured using a liquid scintillation counter, and a fraction of the solubilized product was kept to measure total protein by the Bradford assay. Sample counts were individually normalized to protein, and an average for each transfection was determined. Results were analyzed by paired *t*-test, and statistical significance was determined by p<0.05

### 4.8 RNA Sequencing

INS-1 cells were transfected with either hsTCF19-pcDNA4 or pcDNA4 control vector as stated in the methods above. Cells were cultured 48 hours post-transfection before being collected for RNA using the RNeasy Kit (Qiagen). Total RNA was verified for concentration and purity using a NanoDrop ND-2000c Spectrophotometer and Agilent 2100 BioAnalyzer. Samples that met the Illumina TruSeq Stranded Total RNA (Human/Mouse/Rat) (Illumina Inc., San Diego, CA,USA) sample input guidelines were prepared according to the kits protocol. Cytoplasmic ribosomal RNA reduction of each sample was accomplished by using complementary DNA probe sequences attached to paramagnetic beads. Subsequently, each mRNA sample is fragmented using divalent cations under elevated temperature, and purified with Agencourt RNA Clean Beads (Beckman Coulter, Pasadena, CA, USA). First strand cDNA synthesis is performed using SuperScript II Reverse Transcriptase (Invitrogen, Carlsbad, CA, USA) and random primers. Second strand cDNAs are synthesized using DNA Polymerase I and RNAse H for removal of mRNA. Double-stranded cDNA is purified using Agencourt AMPure XP beads (Beckman Coulter, Pasadena, CA, USA). cDNAs are end-repaired by T4 DNA polymerase and Klenow DNA Polymerase, and phosphorylated by T4 polynucleotide kinase. The blunt ended cDNA is purified using Agencourt AMPure XP beads. The cDNA products are incubated with Klenow DNA Polymerase to add an ‘A’ base (Adenine) to the 3’ end of the blunt phosphorylated DNA fragments and then purified using Agencourt AMPure XP beads. DNA fragments are ligated to Illumina adapters, which have a single ‘T’ base (Thymine) overhang at their 3’end. The adapter-ligated products are purified using Agencourt AMPure XP beads. Adapter ligated DNA is amplified in a Linker Mediated PCR reaction (LM-PCR) for 12 cycles using Phusion™ DNA Polymerase and Illumina’s PE genomic DNA primer set followed by purification using Agencourt AMPure XP beads. Quality and quantity of finished libraries are assessed using an Agilent DNA1000 series chip assay (Agilent Technologies, Santa Clara, CA, USA) and Invitrogen Qubit HS cDNA Kit (Invitrogen, Carlsbad, CA, USA), respectively. Libraries were standardized to 2nM. Cluster generation was performed using the Illumina cBot. Paired-end, 100bp sequencing was performed, using standard SBS chemistry on an Illumina HiSeq2500 sequencer. Images were analyzed using the standard Illumina Pipeline, version 1.8.2. RNA Library preparation and RNA Sequencing was performed by the University of Wisconsin-Madison Biotechnology Center.

Sequencing reads were adapter and quality trimmed using the Skewer trimming program^59^. Quality reads were subsequently aligned to the annotated reference genome (Rnor_6.0) using the STAR aligner ^60^. Quantification of expression for each gene was calculated by RSEM ^61^. The expected read counts from RSEM were filtered for low/empty values and used for differential gene expression analysis using EdgeR^62^ using a false discovery rate (FDR) cut off of < 0.05. Of the 160 genes differentially expressed between the groups, 136 genes were upregulated and 24 were downregulated. Statistical Overrepresentation test and GO term enrichment was performed using PANTHER database on the upregulated genes with fold change >1.5 (http://www.pantherdb.org). Search Tool for the Retrieval of Interacting Genes/Proteins (STRING) database was used to construct the PPI network (https://string-db.org/) ^63^.

### 4.9 Mining Algorithm for GenetIc Controllers (MAGIC) analysis

MAGIC analysis uses Encyclopedia of DNA Elements (ENCODE) ChIPseq data to look for statistical enrichment of transcription factors (TFs) that are predicted to bind to regions in a gene set. It determines if genes in a list are associated with higher ChIP values than expected by chance for a given transcription factor or cofactor based on ENCODE data. All genes that were induced more than 2-fold with an associated FDR < 5% were used as input and tested against the 5Kb_Gene.mtx matrix.

## Author Contributions

GHY, DAF, and DBD conceived and designed the study, GHY, DAF, SL, JAB, AR and DBD completed acquisition, analysis, or interpretation of the data, AR created software used in the work, GHY and DBD drafted or substantially revised the manuscript. All authors approved the submitted version of the manuscript.

## Acknowledgements

We thank Dr. Matthew Flowers for critical review of this manuscript. The authors utilized the University of Wisconsin _ Madison Biotechnology Center’s Gene Expression Center Core Facility (Research Resource Identifier - RRID:SCR_017757) for RNA library preparation, the DNA Sequencing Facility (RRID:SCR_017759) for sequencing, and the Bioinformatics Resource Center (RRID:SCR_017799) for RNASeq quality control and analysis. This work was performed with facilities and resources from the William S. Middleton Memorial Veterans Hospital. This work does not represent the views of the Department of Veterans Affairs or the United States government.

## Data Availability Statement

The Python script and information on how to access required Matrix files are available at https://github.com/asroopra/MAGIC

---

**Supplementary Table 1:**
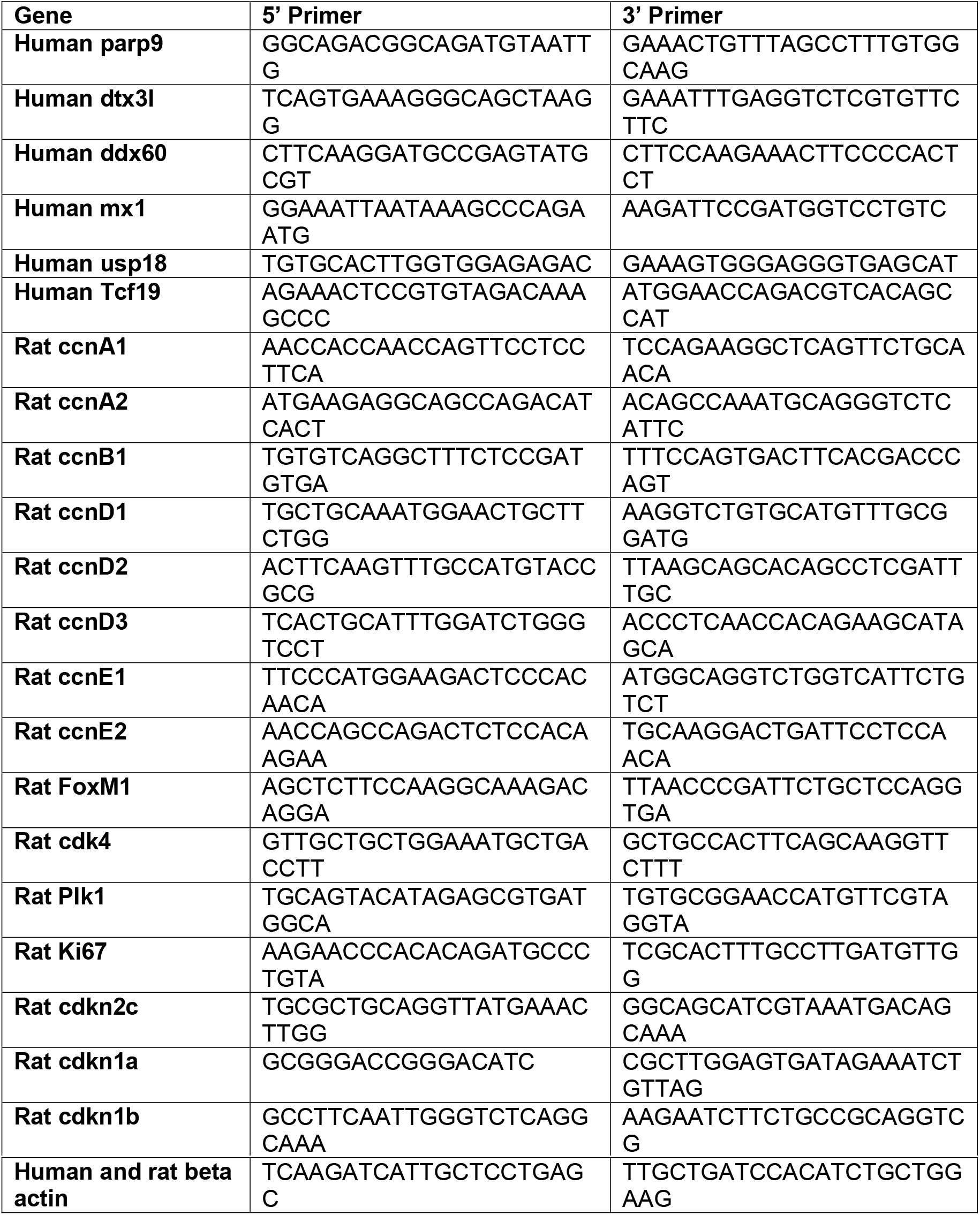
Primer sequences used for real-time PCR experiments.

**Supplemental Table 1**

Differentially expressed genes with FDR<5% from RNAseq on hsTCF19 overexpression in INS1 cells.

**Supplementary Table 2:** Primer sequences used for real-time PCR experiments.

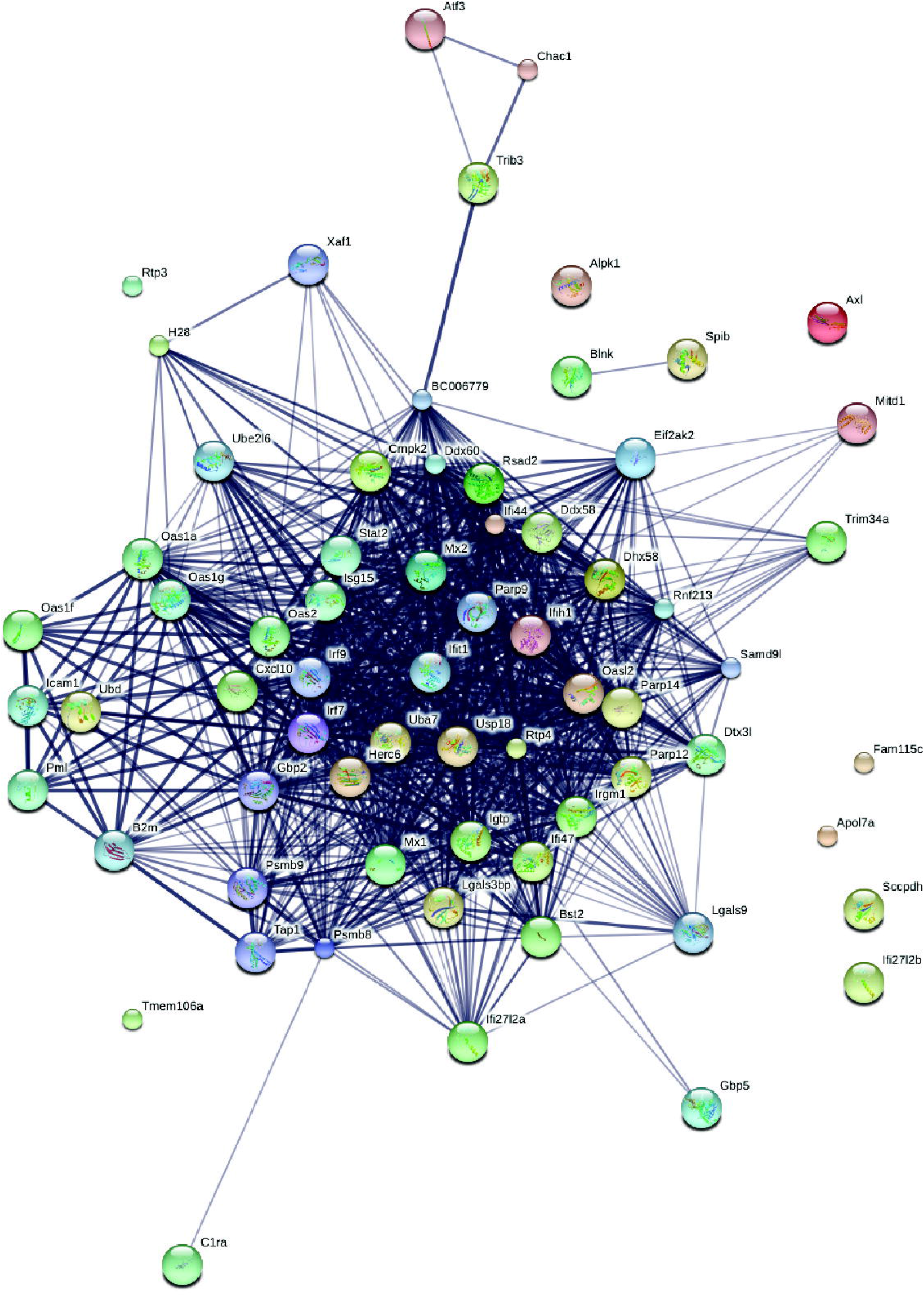

